# Attomolar Detection of HIV-1 with Label-Free RCA-rCRISPR on Smartphone

**DOI:** 10.1101/2025.06.15.659809

**Authors:** Noor Mohammad, Anastasiia Steksova, Yuyang Tang, Leyao Huang, Alireza Velayati, Shengwei Zhang, Aditi Dey Poonam, Sina Jamalzadegan, Matthew Breen, Guochun Jiang, Qingshan Wei

## Abstract

Human Immunodeficiency Virus-1 (HIV) remains a major global public health challenge, having led to over 42.3 million deaths since its discovery in the early 1980s. Despite progress in prevention and treatment, around 60% of people with HIV (PWH) remain undiagnosed in resource-limited regions, disproportionately affecting vulnerable populations and underserved communities across the world. This illustrates the critical need for accessible, accurate, and equipment-free diagnostic tools to enhance detection and thus provide opportunities to curb its spread. Here, we developed a low-cost, robust, and label-free rolling circle amplification (RCA)-rCRISPR diagnostic platform for detecting HIV viral load with minimal instrumentation. Our strategy, combining the integration of RNA-detecting RCA reaction with plasmid reporter-based ratiometric CRISPR (rCRISPR), enables sensitive detection of unprocessed RNA targets without the need for intensive sample pre-treatment. This label-free RCA-rCRISPR diagnostic platform detected HIV RNA down to single-digit aM sensitivity (~3000 copies/mL) from PWH-derived HIV samples *ex vivo*. Unlike typical RCA, which requires sample fragmentations to break long RNA target sequences, our design harnesses the triple functions of the phi29 DNA polymerase (namely exonuclease activity, polymerization, and strand displacement), enabling the detection of the entire HIV genome without pre-fragmentation. For point-of-care (POC) applications, we constructed an all-in-one smartphone-based minigel electrophoresis device to facilitate equipment-free HIV viral load testing, making it accessible to resource-limited communities. Additionally, the assay has demonstrated the ability for point mutation detection (*BRAF* mutation in canine urothelial carcinoma), showcasing the robustness of our strategy for broad disease diagnostic applications.

## Introduction

Human Immunodeficiency Virus-1 (HIV) remains one of the most significant public health challenges globally, with approximately 40 million people living with the virus worldwide as of 2023^1^. Since its discovery in the early 1980s, HIV has led to over 42.3 million deaths^1^, making it a leading cause of morbidity and mortality worldwide. Despite substantial progress in prevention and treatment efforts, around 60% of people with HIV (PWH) are going undiagnosed in resource-limited regions^2^, which means resulting disease disproportionately affects vulnerable populations and underserved communities, highlighting the urgent need for accessible and accurate point-of-care (POC) diagnostic tools to combat the spread of the virus.

The traditional HIV diagnosis relies on the use of antigen/antibody or polymerase chain reaction (PCR) testing^3^. Antibody/antigen tests suffer from low specificity and lack the sensitivity required for reliable early detection, whereas PCR is costly, time-consuming, complex, laborious, and requires a specialized laboratory and skilled technical staff. The technological advancement of gene-editing using CRISPR (<u>C</u>lustered <u>R</u>egularly <u>I</u>nterspaced <u>S</u>hort <u>P</u>alindromic <u>R</u>epeats) with associated proteins (aka. CRISPR-Cas)^4–7^ offers promising opportunities for the development of rapid, sensitive, and specific molecular diagnostics^8–12^. CRISPR-based diagnostics (CRISPR-Dx) came into the limelight after the discovery of Cas13a-based Specific High-Sensitivity Enzymatic Reporter UnLOCKing (SHERLOCK) platform for detecting RNA sequences^9^, and Cas12a-based DNA endonuclease-targeted CRISPR *trans* reporter (DETECTR) mechanism to detect DNA targets^8^. Numerous CRISPR-Dx platforms have been developed to date, showcasing the immense potential of CRISPR systems in addressing the longstanding challenge of achieving highly sensitive and specific nucleic acid detection through a comparatively rapid process^13–28^. CRISPR-Cas12a particularly draws extensive attention, given its ease of reconfiguration for detecting a wide range of human, animal, and plant diseases^8,14,17,20,29–32^, including gene mutations^33,34^. For signal acquisition purposes, CRISPR-Cas12a-Dx typically relies on fluorescent, colorimetric, or electrochemical readout^8,9,13,18,24,26,30,33,35–42^. The signal is mainly achieved when CRISPR-Cas12a nonspecifically cleaves the reporter molecules (typically ssDNA) *via* its unique *trans*-cleavage mode and changes the fluorescence^8,9,13,18,26,33,35,37,38,41^, color^30,36,39,40^, or current states^24,42^ of the reporters. We previously demonstrated a simple, inexpensive, and nonoptical CRISPR-Cas12a-based sensing platform to detect DNA targets at sub-picomolar (pM) detection sensitivity using double-stranded lambda (ds λ) DNA^43^, hybrid DNA (i.e., dsDNA backbone plus a 3’ toe)^44^, and ds plasmid DNA^45^ as label-free reporters. These assays quantify target concentrations by measuring reporter size changes. The plasmid reporter is especially appealing for POC applications, due to its combined benefits of cost-effectiveness, rapid reaction kinetics, high detection sensitivity, and ratiometric signal readout through molecular sizing to reduce size errors.

While Cas12a has been shown to recognize DNA targets, recent studies employing techniques such as split crRNA or Split Activator for Highly Accessible RNA Analysis (SAHARA) have demonstrated the detection of RNA targets using CRISPR-Cas12a^32^. However, RNA detection using these methods often suffers from low detection sensitivity. More recently, researchers combined RCA with fluorescently labeled CRISPR-Cas12a, which enabled the detection of miRNA at single-digit femtomolar (fM) levels^31^. While this approach is effective for miRNA detection, detection of an entire unprocessed viral genome, such as HIV, at clinically relevant levels (low attomolar, or aM) remains challenging. Moreover, the development of a low-cost device for assay readout is crucial if such an assay is to offer a significant impact for adoption in resource-limited communities.

Here, we developed a low-cost, robust, and label-free RCA-rCRISPR diagnostic platform coupled with a smartphone readout device for detecting the whole HIV genome derived from PWH *ex vivo* (**Scheme 1**). This innovation combines a plasmid-based ratiometric CRISPR (rCRISPR) strategy with RNA-detecting RCA. Amplicon DNA generated by the RCA step activates Cas12a, which induces trans-nicking of unlabeled supercoiled ΦX174 (5.37 kb)^46^ plasmid DNA reporter, generating a sensitive ratiometric signal (the ratio of supercoiled to circular plasmid). The assay signal is then quantitatively measured by a portable molecular sizing device such as the smartphone-based mini gel system. Our label-free CRISPR diagnostic platform detected HIV viral load as low as approximately 3,000 copies/mL (~5 attomolar) in PWH-derived HIV *ex vivo*, which is approximately 10^3^ times greater sensitivity compared to typical RCA-integrated fluorescent CRISPR methods. In contrast to conventional RCA methods that necessitate sample pretreatment to detect lengthy genomic RNA, our design harnesses the triple functions of phi29 protein (fast exonuclease activity, polymerization, and strand displacement), enabling the detection of the entire HIV genome without pre-fragmentation. For point-of-care (POC) applications, we have developed an integrated minigel electrophoresis device coupled with a smartphone-based fluorescence reader, enabling equipment-free testing of HIV viral load. Furthermore, our assay exhibits versatility by successfully detecting point mutations, such as the *BRAF* mutation associated with canine urothelial cancer^47–50^, highlighting the robustness and broad applicability of our diagnostic strategy across various diseases.

**Scheme 1:**
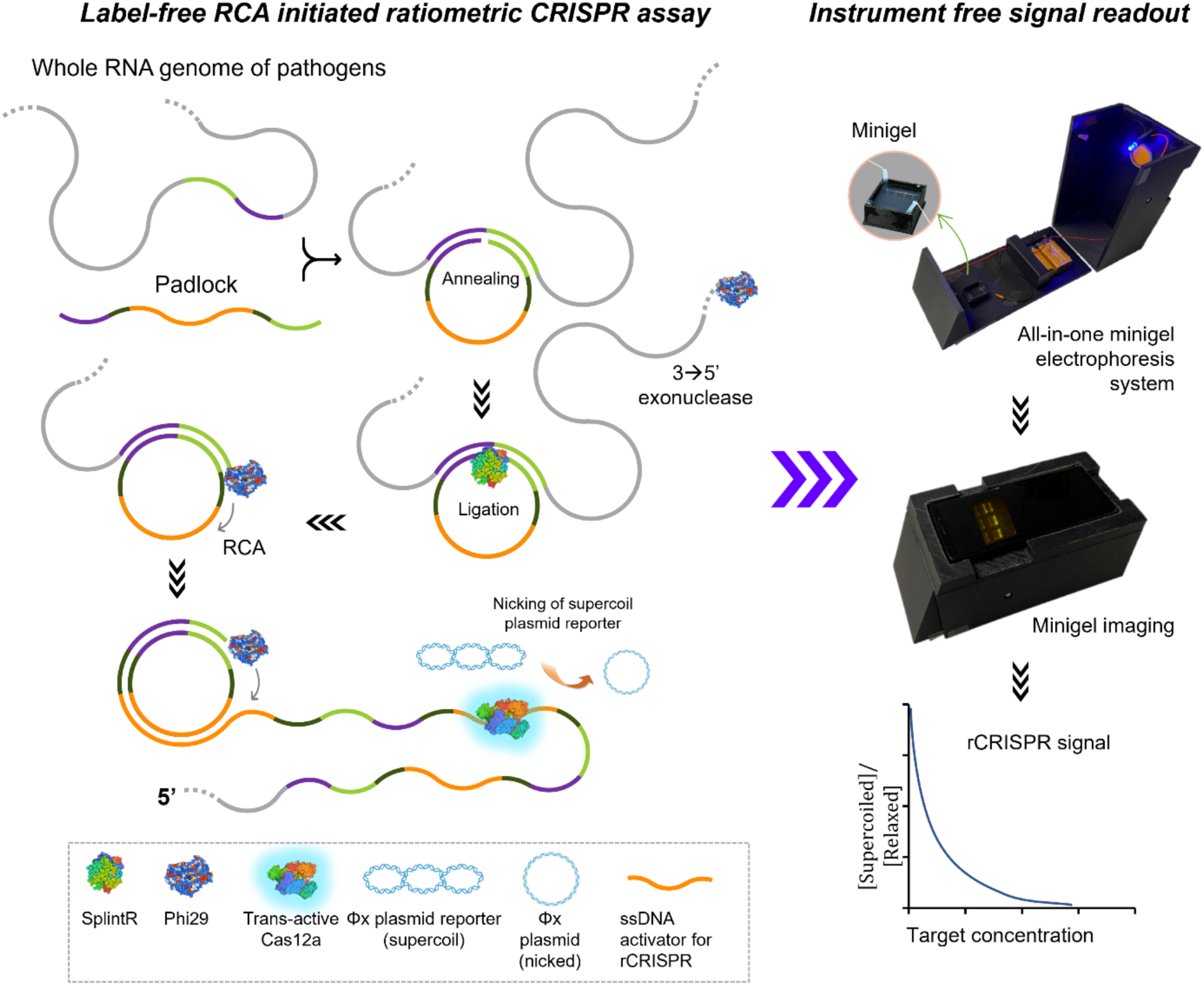
Schematic of plasmid reporter-powered RCA-rCRISPR integrated with a smartphone reader for detecting unprocessed, long HIV genome at POC.

## Results and Discussion

### Integrating Label-free rCRISPR with RCA for RNA detection

rCRISPR is a novel CRISPR-Cas12a-based diagnostic method for detecting DNA targets using a label-free plasmid reporter molecule^51^, and the RCA assay is an isothermal nucleic acid amplification method. The combination of these two allows us to expand the detection capability of rCRISPR from DNA targets to RNA.

We first tested the RCA and rCRISPR assays separately and studied the feasibility of detecting RNA molecules using a label-free plasmid reporter. **Fig. 1a** shows the mechanism of RCA assay. We designed an RCA padlock to detect a 22 nt long synthetic RNA target (sequences of padlock and 22 nt RNA target are in **Table S1**). The 3’-end of the target has reverse complementarity to the 5’ end of the padlock, and the 5’-end of the target has the reverse complementarity to the 3’ end of the padlock. This complementary matching design helps the target nucleic acid bind to both ends of the padlock and join the two ends of the padlock together. The SplintR ligase enzyme then ligates both ends of the padlock and produces a circular template, on which the target RNA acts as the primer for the RCA assay. Next, the phi29 DNA polymerase starts synthesizing DNA in a 5’ to 3’ direction. With a unique ability to displace the downstream DNA strand, phi29 continues DNA synthesis even when encountering secondary structures or other obstacles in the template DNA. In this manner, phi29 keeps rolling over the circular template and produces many copies of reverse complementary sequences of padlock with the long ssDNA product >10 kb. We performed RCA for 2 hrs at room temperature and then ran gel electrophoresis to evaluate the reaction products. **Fig. 1b** (lanes 3 and 4) showed that in the presence of RNA targets, RCA reaction occurred successfully. We observed large molecular weight DNA products (bands) adjacent to the wells, confirming the presence of RCA products. In contrast, in the absence of RNA targets, no amplification was noticed (**Fig. 1b**, lane 2). These results demonstrated that an RNA-detecting RCA assay was successfully constructed, consistent with prior studies^31^.

**Fig. 1:**
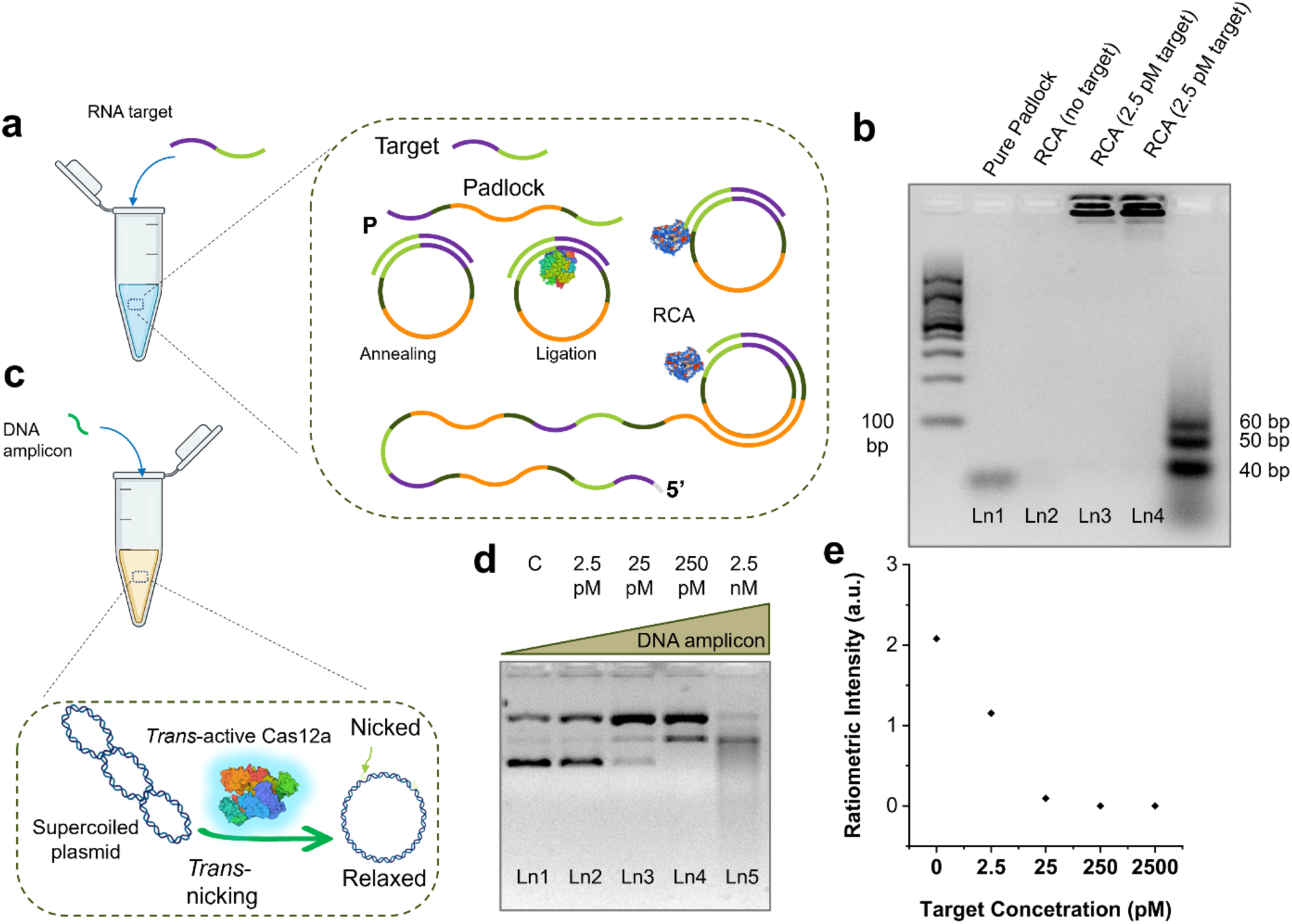
Demonstration of RCA and label-free rCRISPR separately. (a) Schematic of RCA for RNA detection. (b) Gel electrophoresis (4 % agarose gel and 1×TBE buffer) results demonstrating results for RCA assay performed at room temperature for 2 hrs. (c) Schematic demonstration of label-free rCRISPR for ssDNA detection using ΦX174 plasmid reporter. (d) Gel electrophoresis (1% agarose gel and 1×TBE buffer) results demonstrating rCRISPR assay performed at 37 °C for 1 hr. Ln1: negative control; Ln2-5: increasing concentrations of target. (e) Ratiometric signal achieved from rCRISPR strategy. Abbreviations: RCA, Rolling circle amplification; c, negative control; nM, nanomolar; pM, picomolar; ssDNA, single-stranded DNA; bp, base pair; Ln, lane; TBE, Tris-borate EDTA; a.u., arbitrary unit.

Next, we confirmed the feasibility of detecting ssDNA using rCRISPR with a ds plasmid reporter (supercoiled ΦX174). To demonstrate the rCRISPR assay with ΦX174 supercoiled plasmid reporter, we first activated the Cas12a with a 20 nt long ssDNA (see sequence in **Table S1**), and added plasmid reporter in the reaction. **Fig. 1c** shows the schematics of label-free CRISPR-Cas12a assay using a supercoiled plasmid reporter. We performed the assay with 2.5 pM, 25 pM, 250 pM, 2.5 nM of targets, with a negative control. After 1 hr of reaction, the gel electrophoresis showed that supercoiled bands became thinner for all of the concentrations except for the negative control (**Fig. 1d**). We measured band intensities (**Fig. S1**), and plotted ratiometric signals (ratio of supercoil to circular band) against different target concentrations. **Fig. 1e** shows that with the increase of target concentrations, the ratiometric signal went down and became zero for higher target concentrations above 25 pM.

Then, we combined both assays together to detect RNA targets (**Fig. 2a**). We mixed all the required reagents for RCA and rCRISPR in a single pot and incubated the reaction mixture for 2 hrs at 37 °C. We performed the assay with RNA targets of 250 pM, 2.5 nM, 25 nM concentrations, and 0 target (negative control) to evaluate the feasibility of RCA-rCRISPR scheme for RNA detection. **Fig. S2a** shows that in the initial trial, we didn’t observe any supercoil relaxation for any of the target concentrations. After careful troubleshooting of all assay conditions, we attributed the failure of the direct combination of RCA and rCRISPR reactions to the opposite functionality of SplintR ligase and Cas12a enzyme. We suspected that while target*-*activated Cas12a tended to nick and unwind supercoiled plasmid reporters, the SplintR ligase on the other side would repair the Cas12a-induced nicks and recover its supercoiled status (**Fig. S2b)**. One potential way to circumvent this problem is to deactivate the SplintR ligase before the subsequent rCRISPR reaction. To test that, we first ran the RCA reaction at room temperature and then deactivated the ligase protein at 65 °C before performing the rCRISPR at 37 °C. The procedure of the assay and gel results are shown in **Figs. 2b and c**, respectively. The gel results showed a positive rCRISPR assay result this time, confirming that Cas12a-induced DNA supercoil relaxation happened after thermal denaturation of RCA ligase.

**Fig. 2:**
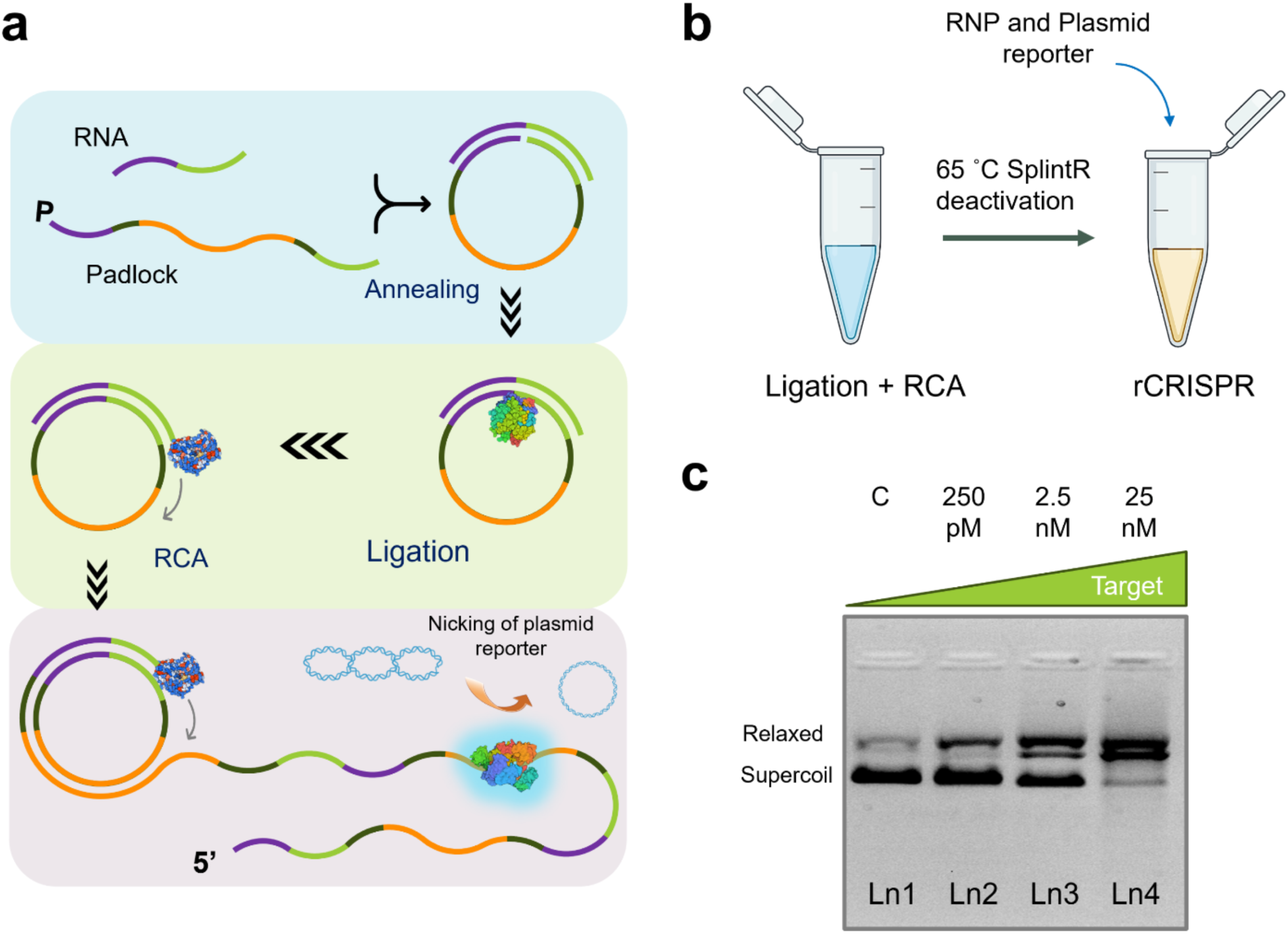
RNA detection using RCA-rCRISPR. (a) Mechanism of RCA-integrated rCRISPR. (b) Schematic of SplintR denaturation. (c) Gel electrophoresis (1% agarose gel and 1×TBE buffer) results demonstrating RNA detection using RCA-rCRISPR assay. RCA, Rolling circle amplification; c, negative control; nM, nanomolar; pM, picomolar; Ln, lane; TBE, Tris-borate EDTA; dsDNA.

To simplify the assay protocols and explore the possibility of combining two separate assays in one pot, instead of deactivating SplintR, as ATP is an essential molecule for ligation assay, and can be degraded at temperatures in excess of 40 °C^52,53^, we chose to inactivate ATP once ligation is complete. On the other side, rCRISPR can be performed at higher temperatures (>45 °C). The idea is to perform rCRISPR at an ATP-susceptible temperature (e.g., 52 °C) to inhibit plasmid reporter repairment during the CRISPR reaction. To test the idea, we designed three different assay protocols: i) two-step RCA-rCRISPR with SplintR deactivation at 65 °C, ii) two-step RCA-rCRISPR at 52°C without SplintR deactivation step, and iii) one pot RCA-rCRISPR assay at 52°C (**Figs. 3a, d, and g**).

**Fig. 3:**
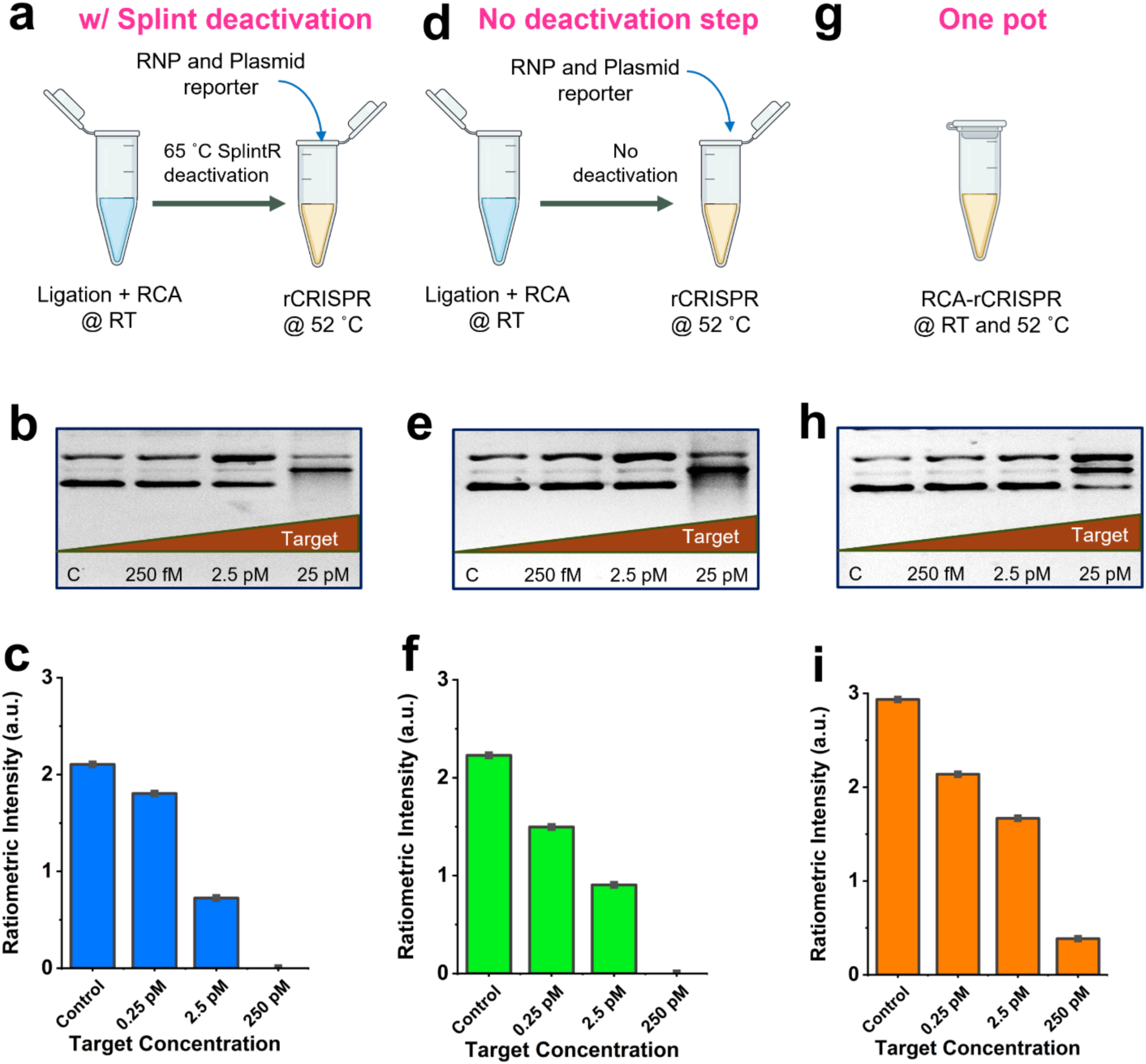
Assay simplification. Schematics show experimental procedures with ligase deactivation (a), without ligase deactivation (d), and one pot assay (g). Gel electrophoresis (1% agarose gel and 1×TBE buffer) results demonstrating RNA detection using rCRISPR with ligase deactivation (protocol i) (b), without ligase deactivation (protocol ii) (e), and one pot assay (protocol iii) (h). Histogram showing quantitative ratiometric signals for the assays with ligase deactivation (protocol i) (c), without ligase deactivation (protocol ii) (f), and one pot assay (protocol iii) (i). Abbreviations: RCA, Rolling circle amplification; RT, room temperature; c, negative control; pM, picomolar; Ln, lane; TBE, Tris-borate EDTA; a.u., arbitrary unit.

For protocol i, the assay worked as expected (**Fig. 3a-c**). For protocol ii (without SplintR deactivation), the RCA assay tubes (without CRISPR reagent) were kept at RT for 2 hrs and then CRISPR reagents were added. The tubes were incubated at 52°C for 1 hr. For protocol iii (one-pot assay), we combined everything in one tube kept it at RT for 2 hrs and then moved it to an incubator at 52°C. The concept is that the ligation reaction would be favorable at a lower temperature^54^. At a higher temperature, ATP will be degraded, thus inhibiting the ligation reaction^53^ and leading to a positive rCRISPR signal. Qualitative gel results (**Figs. 3b, e, and h**) and quantitative ratiometric signal measurement (**Figs. 3c, f, and i**) showed the success of all three assay protocols. However, if we compare the supercoil bands for the 250 pM targets in **Figs. 3b, and e**, we noticed that protocol ii (without deactivation) generated the strongest signal. Therefore, we followed protocol ii for all future experiments. We also evaluated the effect of padlock concentrations on the final assay signal intensity (**Fig. S3**) and found that the higher padlock concentration (250 nM) hampered the overall assay signals. This might be due to the binding of two padlocks to one target when the padlock concentration is too high. This side effect would reduce the formation of the desirable circular templates for RCA and therefore reduce signal intensity (**Fig. S4**).

### HIV detection using the RCA-rCRISPR technique

After optimizing and simplifying the assay, we tested the new RCA-rCRISPR assay for HIV target detection. We chose the pol region of the HIV genome as our target (**Fig. 4a**, also see sequence in **Table S1**)^55^. An RCA padlock was designed to recognize a 22-nt sequence from the pol region (see the sequence in **Fig. 4a**). For the proof-of-concept demonstration, we synthesized a 28-nt (3 nt extra on either side) sequence to mimic the target of the HIV genome. The extra nucleotides (nts) were added to evaluate the assay performance when the target was not the exact size of the hybridization site of the padlock.

**Fig. 4:**
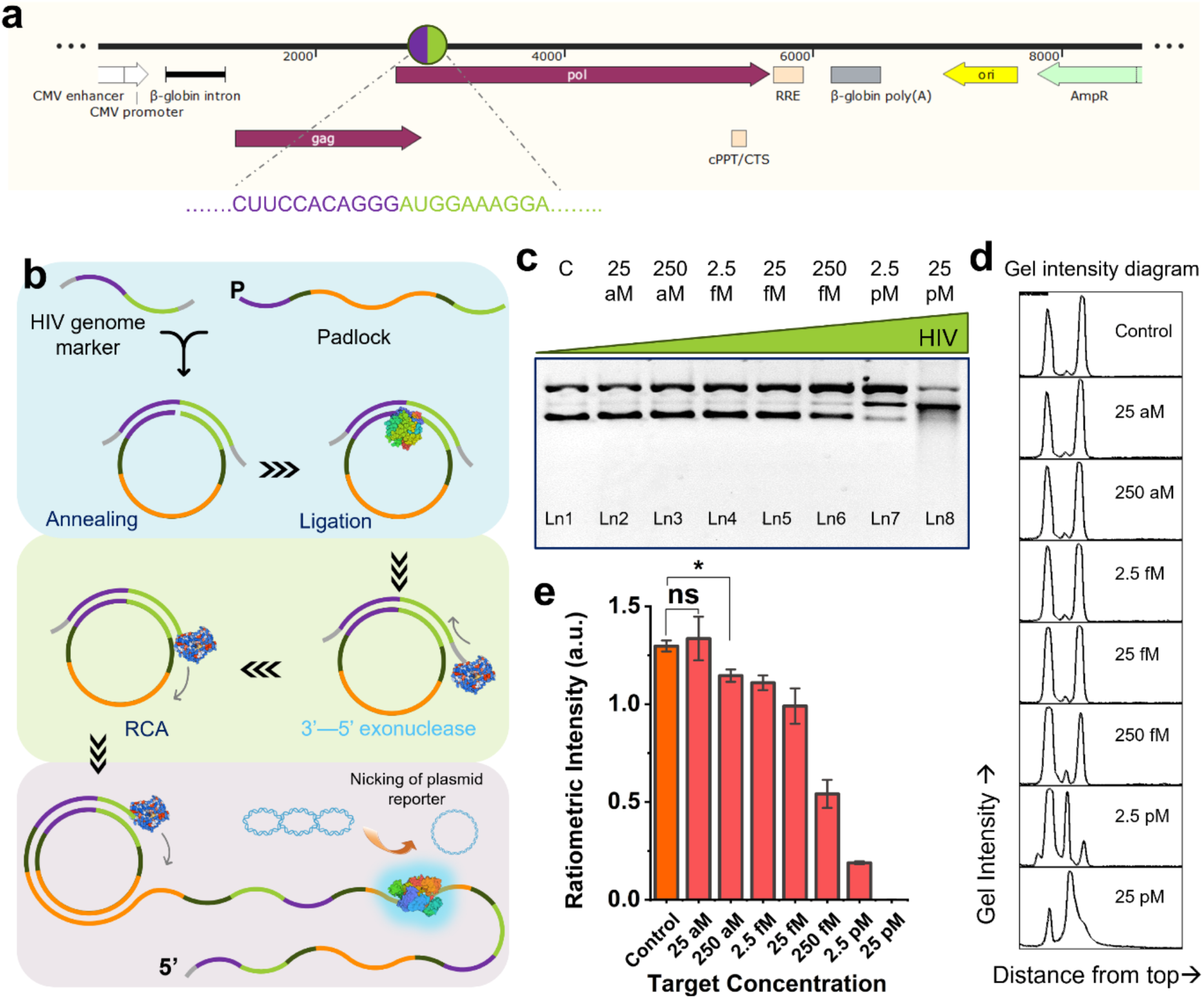
Synthetic HIV RNA (pol gene) detection using RCA-rCRISPR technique. (a) HIV RNA target selection. (b) Assay mechanism of HIV RNA detection utilizing RCA-initiated rCRISPR technique. (c) Gel electrophoresis (1% agarose gel and 1×TBE buffer) demonstrating rCRISPR assay result for HIV RNA target detection. (d) Gel intensity diagrams of each lane in c. (e) Quantitative performance (LOD quantification for synthetic HIV RNA) of rCRISPR. The graph shows statistical insignificance at p>0.05 (ns), and significance at p<0.05 (*). Abbreviations: c, negative control; aM, attomolar; fM, femtomolar; pM, picomolar; Ln, lane; TBE, Tris-borate EDTA; a.u., arbitrary unit.

**Fig. 4b** shows the mechanism of detecting the 28 nt synthetic HIV target using the RCA-rCRISPR strategy. The main difference in this schematic compared to the previous one in **Fig. 2a** is the exonuclease activity of phi29. Here, since we have three extra nts at the 3’ end of the target, we propose that when polymerization starts, these extra nucleotides will be removed by phi29. After that, the HIV target would act as the primer for RCA amplification. We performed the assay for a wide range of target RNA concentrations, ranging from 25 aM to 25 pM, including a negative control (**Figs. 4c and d**). To evaluate the assay performance, we measured the limit of detection (LOD), which was found to be ~250 aM.

We next applied the optimized assay to measure HIV RNA in samples directly isolated from PWH. We obtained HIV samples, which were originally isolated from the brain tissue of PWH, from the HIV Cure Center at the University of North Carolina at Chapel Hill (UNC-Chapel Hill). The donor was undergoing antiretroviral therapy (ART) and had an undetectable plasma viral load^56^. Therefore, instead of attempting direct detection, we used PWH-derived HIV samples to perform *ex vivo* infection of primary brain microglia. Our data showed that we were able to detect the viral RNA release in culture supernatants in this *ex vivo* infection study. **Fig. 5a** demonstrates the steps involved in the detection of PWH-derived HIV RNA using label-free RCA-rCRISPR. Briefly, the collected viruses from the cultured supernatants were lysed in 1% Triton X-100 to release their viral load. Viral RNA was purified using a spin column and then added to the label-free RCA-rCRISPR assay. **Fig. 5b** shows the 3’→5’ exonuclease activity of phi29 to trim the long whole genome target and convert HIV RNA into the primer for further RCA amplification. **Fig. 5c** shows the assay result with the whole HIV genome from the PWH-derived HIV samples. Lane 1 was for control, and lanes 2 and 3 were for the assay with samples containing PWH-derived unprocessed HIV. To confirm that these results are not false positive, we first verified that the viral target sequence we chose did not match the human genome (**Fig. S6a**). In addition, since we know that our label-free assay is based on the *trans-*nicking of the plasmid reporter, we ran assays with and without the padlock to confirm that the supercoil relaxation in **Fig. 5c** is not due to unknown interferences from samples with PWH-derived HIV (**Fig. S6b**).

**Fig. 5:**
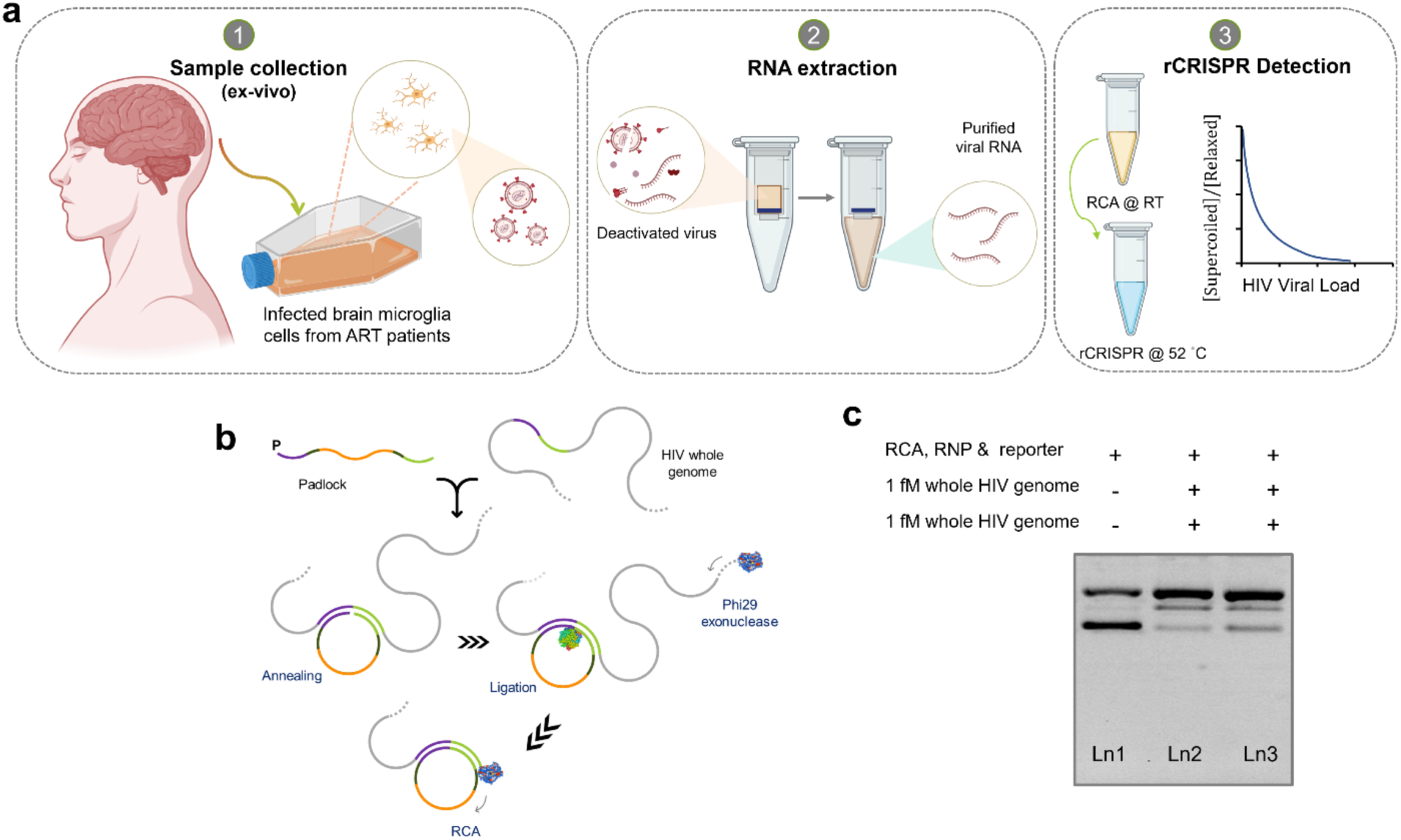
Ex vivo HIV RNA detection with RCA-rCRISPR. (a) Schematic showing the overall procedures of detection of the HIV genome from PWH. (b) Mechanism showing how a very large viral RNA (i.e., whole HIV genome without ultrasonication) can initiate RCA reaction without a primer. (c) Gel electrophoresis (1% agarose gel and 1×TBE buffer) demonstrated rCRISPR assay for whole HIV genome detection. Abbreviations: fM, femtomolar; Ln, lane; TBE, Tris-borate EDTA.

After the initial demonstration of detecting an unprocessed whole HIV genome, we then quantified the limit of detection (LOD) of RCA-rCRISPR for HIV derived from PWH. We serially diluted the samples in nuclease-free water and ran the assay with 0.5 aM, 5 aM, 50 aM, 250 aM target RNA concentrations with a negative control (**Fig. 6**). We ran gel electrophoresis to observe the supercoil relaxation of the plasmid reporters (**Fig. 6a**). Then, we determined the LOD by measuring the ratiometric band intensity (**Fig. 6b**). We observed that HIV viral load can be detected at a single digit aM level (5 aM, **Fig. 6c**). It is worth mentioning that the LOD for whole HIV genome was much better than the shorter synthetic targets (**Fig. 4e**). We hypothesized that for longer targets, the 3’→5’ exonuclease activity of phi29 might be higher, which requires further investigation.

**Fig. 6.**
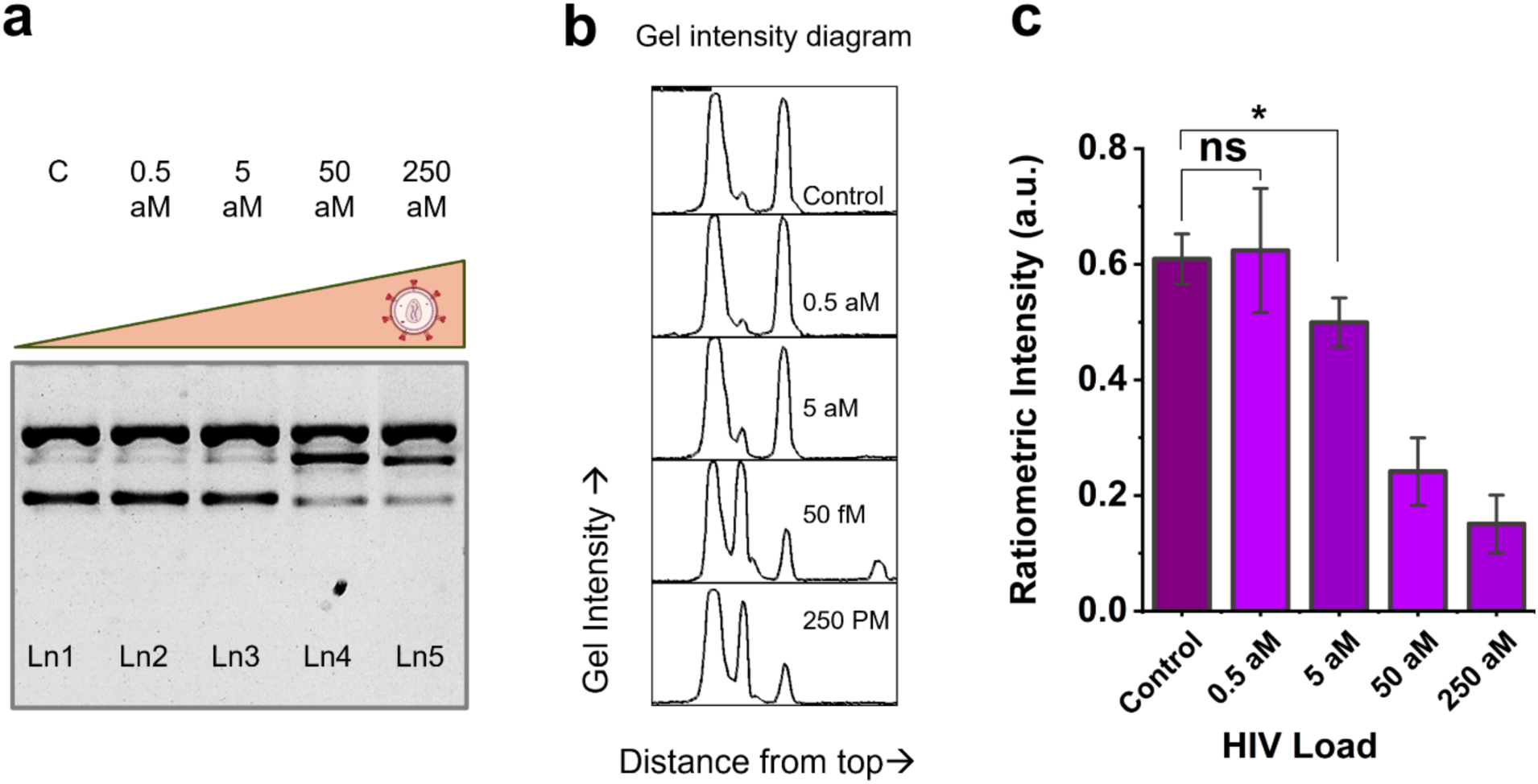
LOD quantification for the whole HIV genome detection using RCA-rCRISPR strategy. (a) Gel electrophoresis (1% agarose gel and 1×TBE buffer) results demonstrating RCA-rCRISPR assay result for the whole HIV genome detection. (b) Gel intensity diagrams of each lane in a. (c) LOD quantification for HIV whole genome derived from PWH. The graph shows statistical insignificance at p>0.05 (ns), and significance at p<0.05 (*). Abbreviations: c, negative control; aM, attomolar; Ln, lane; TBE, Tris-borate EDTA; a.u., arbitrary unit.

If we compared our results with previously demonstrated typical fluorescent reporting model that utilizes RCA-rCRISPR-Cas12a for RNA detection, we found that our low-cost label-free ratiometric reporting strategy performed three orders of magnitude (10^3^ times) better than the widely labeled F-Q reporting system^31^. Further, this ratiometric sensing strategy is advantageous from additional perspectives. Notably, DNA supercoil relaxation is rapid and can easily be evaluated using a more compact miniaturized gel electrophoresis system^57,58^ instead of a benchtop fluorescence microscope for POC detection, which is illustrated in the next section.

### Development of a smartphone-supported mini gel electrophoresis system for POC RCA-rCRISPR

Around 60% of PWH remain undiagnosed in resource-limited, underserved communities. Therefore, it is crucial to develop an equipment-free diagnostic system that can be deployed to such regions. The benchtop gel electrophoresis system requires an external power source, an expensive gel imager, and a computer to acquire gel images and analyze the data. Also, reagent consumption is high, which limits this strategy for POC use. Few previous studies demonstrated that microfluidic gel electrophoresis was feasible, but each system came with its own limitations. For instance, use of polymer-based^59^, glass-based^60^, and 3D printed^61^ microfluidic gel electrophoresis have been reported, but these setups are for single-lane gel and require high voltages (70-200V). Although a low-voltage microelectronic-based gel electrophoresis^62^ has been reported, the chip fabrication is very complicated.

We developed an all-in-one minigel electrophoresis system that requires a very low amount of power. This minigel system comprises a minigel cassette, a power supply (9V batteries), one blue LED (Amazon, ~3.21 lumens) to excite the gel during imaging, a fluorescence emission filter (1-inch diameter, 550-nm cutoff wavelength), and a smartphone holder for smartphone integration. To construct the minigel, a rectangular tank (2 cm × 2.5 cm × 0.75 cm) was printed using a 3D printer (Stratasys UPrint SE Plus). We installed two pieces of thin platinum wire (0.25 mm diameter) at the two opposite ends of the tank to make conductive electrodes. These electrodes were then connected to the battery using conductive wire. A blue LED was also connected to the same battery as the excitation light source. All construction steps, electrophoresis feasibility, preliminary demonstration, and system optimization steps have been shown in (**Figs. S7-S10**). This minigel device requires no external power, reduces the reagent consumption massively, and enables naked eye/smartphone detection.

We tested the performance of the smartphone-minigel system to run the label-free RCA-rCRISPR for the detection of PWH-derived HIV genomes (**Fig. 7**). In the minigel device, 2 µL of RCA-rCRISPR assay product was mixed with 1X loading dye and added into the holes in the minigel. After the completion of gel electrophoresis, a smartphone was placed on the top of the device to capture the gel images under blue LED illumination. 5 aM, 50 aM target concentrations, and a negative control were tested and compared. We observed that the supercoil band for 50 aM was significantly different (thinner) than the control band, meaning that our smartphone-based minigel device was able to detect a very low amount of viral load with minimal instrumentation.

**Fig. 7:**
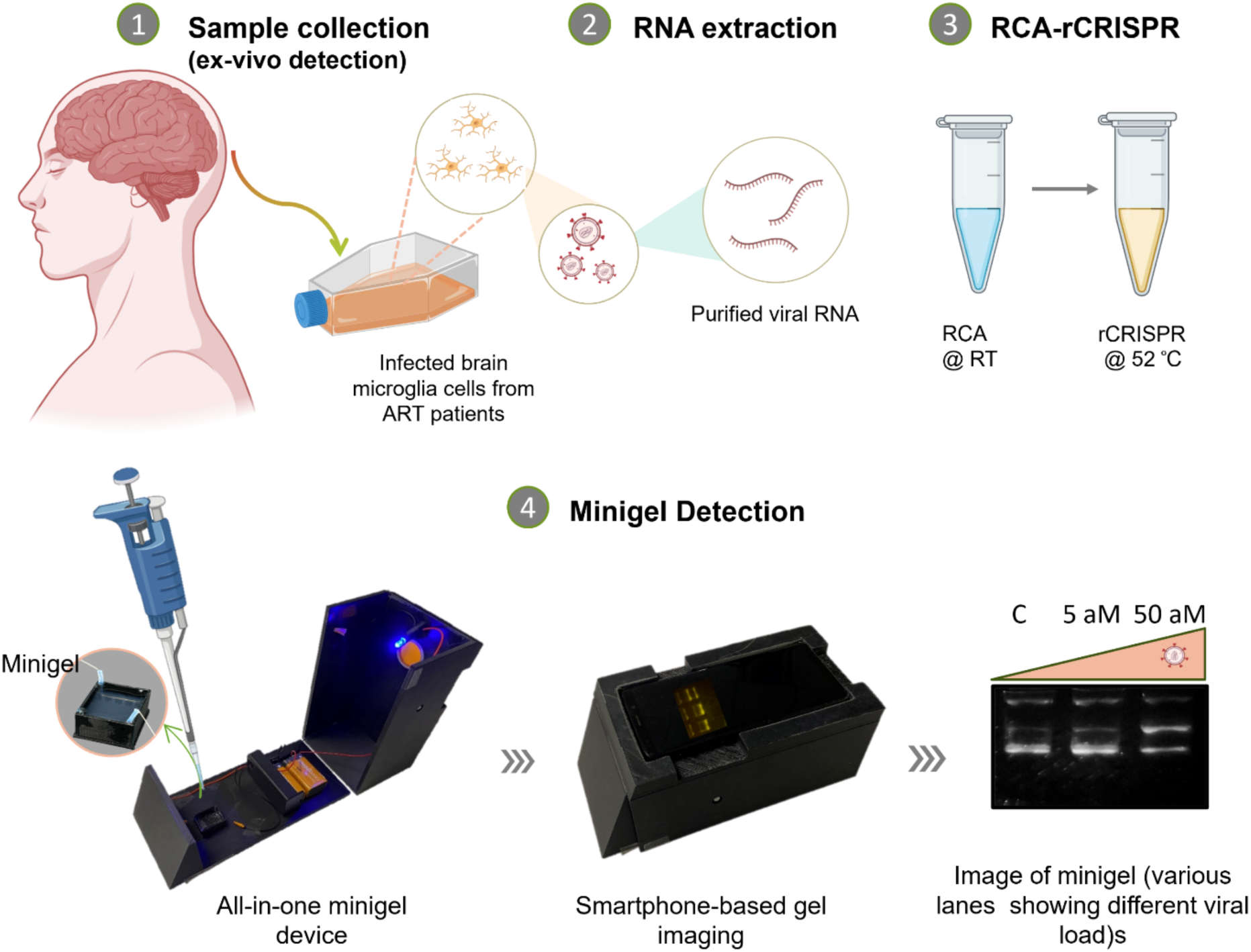
Detection of PWH-derived HIV RNA at POC using the smartphone-minigel system. Abbreviations: c, negative control; aM, attomolar.

### Point mutation detection with RCA-rCRISPR

The RCA method is also known as a powerful technique that can be used to detect single nucleotide polymorphisms (SNPs) in DNA. As a model SNP target, we considered the *BRAF* gene that encodes a protein called B-Raf. In humans, mutations in the *BRAF* gene are associated with various cancers, particularly melanoma, colorectal cancer, and thyroid cancer. One of the most common mutations in the *BRAF* gene is the V600E mutation, where a valine (V) is substituted with glutamic acid (E) at position 600 of the protein. This mutation results in the constitutive activation of the *BRAF* protein kinase, leading to uncontrolled cell growth and proliferation, which is a hallmark of cancer^48^. In the domestic dog, the same base change, referred to as V595E, has a remarkably high frequency (85%) in canine urothelial carcinoma^47,48^. We therefore chose to assess the use of our label-free RCA-rCRISPR strategy for sensitive detection of this single base change in the *BRAF* gene. In the presence of wild-type target, the 5′-terminal with phosphate (p) and 3′-terminal of the padlock probe join together by SplintR DNA ligase when the base at the 3′-end of the padlock probe is perfectly matched to the DNA target. However, this circularization is hampered if there is a mismatch at the joining point. After that, the ligated padlock probe, the extended 3′-end of the target gets trimmed with phi29 protein. Once the exonuclease activity is completed, the remaining sequence of the target acts as the primer for RCA amplification. The RCA amplification produced lots of amplicons, which further activated the CRISPR-Cas12a system and resulted in the relaxation of supercoiled plasmid DNA reporters. This overall detection mechanism has been demonstrated in **Fig. S12a**. The assay was performed with 25 fM of synthetic mutated DNA and wild-type DNA (**Fig. S12b** and **c**). In **Fig. S12c**, we observed that while the supercoil band for the mutated DNA became significantly thinner (lane 3), the same band for wild type (lane 2) was comparable to the control band (lane 1). This result verifies the robustness of our assay platform for point mutation detection.

This study developed a new RNA detection method using a label-free ratiometric CRISPR-Cas12a technique using a recently discovered plasmid reporter. We successfully detected unprocessed whole genome RNA with a detection sensitivity 10^3^ times greater than previously demonstrated fluorescent RCA-CRISPR-Cas12a technique. A portable, battery-powered, smartphone-based minigel electrophoresis system was also developed to quantify ratiometric signals, enabling on-site testing without sophisticated equipment. Compared to lateral flow CRISPR assays^18,63^ and microfluidic digital CRISPR assays^27,38^, the all-in-one gel electrophoresis device for CRISPR detection is much easier to scale up and implement at the POC. This innovative, low-cost, and minimally instrumentalized approach offers a promising solution for combating the ongoing HIV pandemic in resource-limited settings.

However, several challenges remain for the further development of the POC detection system. For instance, sample preparation (extracting HIV viral RNA) still relies on lab-centered techniques (e.g., commercial nucleic acid extraction methods). To simplify the nucleic acid extraction step, nuclease deactivation and heat treatment could be incorporated in future detection workflow^40,64^. Additionally, detecting HIV in minimally equipped areas also poses regulatory challenges, because handling HIV samples typically requires a BSL-2+/BSL-3 laboratory^65^, which would increase the cost and limit the accessibility. In that respect, better and safer sample collection and storage protocols or methods will be needed to collaborate with the new diagnostic approaches.

## Conclusions

In summary, this study demonstrated a POC friendly RCA-rCRISPR system for sensitive HIV RNA detection, with four unique and useful aspects: First, it shows plasmid reporter can be combined with powerful CRISPR reaction schemes (e.g., RCA-rCRISPR-Cas12a) for label-free RNA detection. Second, the molecular sizing-based RCA-rCRISPR assay was 1000 times more sensitive than those previously demonstrated with fluorescent RCA-CRISPR systems, which detected HIV down to ~3,000 copies/mL (5 aM). Considering that the HIV viral load ranges from 15,000 to 50,000-200,000 copies/mL^66^ during acute and chronic infection in PWH, our RCA-rCRISPR detection of HIV is sensitive enough to capture active HIV infection in clinics. Third, unlike conventional RCA-CRISPR, where ultrasonication or additional priming is needed for detecting larger targets, we demonstrated the detection of the whole HIV genome without any nucleic acid fragmentation step, which simplified the assay. Fourth, a battery-powered, smartphone-based all-in-one minigel electrophoresis system was also developed to quantify RCA-rCRISPR signals in an equipment-free manner. Together, this straightforward yet sensitive strategy could be the gateway to accelerate the dissemination of POC HIV tests to resource-limited communities, which is direly needed for early detection of HIV infection with a smartphone to prevent the pandemic globally.

## Methods

### Reagents and apparatus

LbaCas12a, crRNA, synthetic RNA targets, and crRNA were purchased from Integrated DNA Technologies (IDT, Coralville, IA, USA). NEBuffer™ r2.1, and ΦX174 DNA (Catalog# FERSD0031) were purchased from Thermo-fisher. Ultrapure water (18.3 MΩ cm) was produced by the Milli-Q system (Millipore, Inc., USA) and used throughout the experiments. The sequences of ssDNA oligonucleotides and crRNA molecules are listed in Table S1.

### Label-free rCRISPR (ratiometric) assays for DNA detection

All DNA and RNA oligos were stored in IDTE buffer (pH 8) at 10 μM concentrations in a −20 °C refrigerator and pre-warmed at 37 °C for 20 min before mixing. Reporter molecules (e.g., ΦX174) were diluted to 50 ng/µL in IDTE buffer (pH 8) and stored at 4°C for use throughout the experiments. For label-free rCRISPR assay, analyte solutions with different concentrations of ssDNA were added into the CRISPR reaction, consisting of 40 nM LbaCas12a, 20 nM gRNA, 4 ng/ µL reporter molecule, and 1X reaction buffer (e.g., 50 mM NaCl, 10 mM Tris-HCl, 10 mM MgCl_2_, 100 µg/ml recombinant albumin, pH=7.9 @ 25°C). The total reaction volume was 40 µL. Gel electrophoresis was performed to observe the change in conformation of the plasmid reporter. 1% agarose gels were prepared using 1× TBE (Tris Borate EDTA) and 1× SYBR™ Safe DNA Gel Stain (Thermo-fisher, catalog# S33102). 10 μL of different reaction products with loading dye (5:1, v/v) were added to each well. Electrophoresis was done at 120 V for 30 min in the same buffer at room temperature. Finally, the agarose gels were scanned and recorded by the E-Gel Imager system (Invitrogen, USA).

### Label-free rCRISPR assays for RNA detection

For RNA detection, three different mixtures were prepared, A: 100 nM padlock mixed with various concentrations of RNA target in 1x SplintR buffer; B: 400 units/mL SplintR ligase (catalog# M0375S), 125 units/mL phi29 polymerase (catalog# M0269S), 0.5 mM dNTPs, and 250 ng/uL recombinant albumin in 1.25X SplintR ligase buffer; C: 225 nM LbCas12a with 112.5 nM crRNA in 1X NEB r2.1 buffer. For one pot assay, A, B, and C were added in in 1:4:1 ratio. Then 4 ng/ µL label-free plasmid reporter was added. For two-pot assay, the same amount of reagents was added only difference is that mixture C was added after a 2-hr incubation of ‘A+B’. Then, gel electrophoresis was performed to evaluate the assay performance.

### Detection of HIV genome derived from PWH

As most HIV-positive individuals in the U.S. are under effective ART and typically have undetectable plasma viral loads, we performed ex vivo infection in primary brain microglia with HIV isolated from PWH (PMID: 37317962, donor 2), who was virally suppressed by ART. Two days post-infection, cultures were extensively washed to remove residual input virus. Supernatants from the infection culture were subsequently collected 7 days post-infection for HIV RNA detection in this study. The virus in the supernatant was inactivated and lysed using 1% Triton X-100, followed by viral RNA extraction using the Zymo Viral RNA Kit (Catalog #R1035), according to the manufacturer’s instructions. RNA was eluted in 15 µL of DNase/RNase-free water.

### Equipment free HIV RNA detection using smartphone readout minigel platform

A piece of precast gel (1% agarose gel and 1× TBE buffer; gel size: ~1×1.5×.5 cm3) was placed in the minigel tank. 2 µL of reaction including 1× loading dye was added to each lane and run for 1 hr in the battery-powered minigel device. After that, a blue LED switch was turned on, and an image was captured using a smartphone (Samsung Galaxy S9).

### Electrophoresis analysis

The gel results underwent analysis using ImageJ software. Initially, the background of the gel was removed. Subsequently, the intensity along the length of each lane was averaged across its width and graphed. Gel intensity was determined by identifying distinct peaks corresponding to supercoiled and relaxed circular conformations of DNA. Ratiometric intensities, representing the ratio of supercoiled DNA to relaxed circular DNA, were derived by dividing the intensity of the supercoiled band by that of the relaxed circular band. Finally, these ratiometric intensities were plotted against the target concentration.

### Statistical analysis

The student’s t-test was conducted using repeated experimental data (n=3) to compute p-values, thereby assessing the distinction between experimental outcomes and control groups and establishing the limit of detection (LOD). The graphical representation illustrates statistical insignificance when p>0.05 (ns) and statistical significance when p<0.05 (*).

## Supporting Information

The supporting information is available free of charge at https://

## Author Contributions

Q.W. and N.M. initiated and conceived the project. N.M. performed all the experiments, collected, and analyzed the data. A.V. and S.Z. designed the cellphone holder. Y.T. and G.J. prepared HIV ex-vivo patient samples. A.S. and L.H. helped run the gel experiments to produce triplicate data. A.D.P. and M.B. provided SNP mutation sequences. S.J. shared synthetic HIV target sequences. N.M. and Q.W. wrote the manuscript. All the authors helped revise the manuscript.

## Acknowledgments

The authors sincerely thank the funding support from the National Science Foundation (Award # 1944167). G.J. is supported by NIMH (R01MH136852) and NIAID (R21 AI167709-01A1 and R01AI186609).

