## Supporting information for "Attomolar Detection of HIV-1 with Label-Free RCA-rCRISPR on Smartphone"

**Nonfluorescent Label-free rCRISPR for RNA detection**

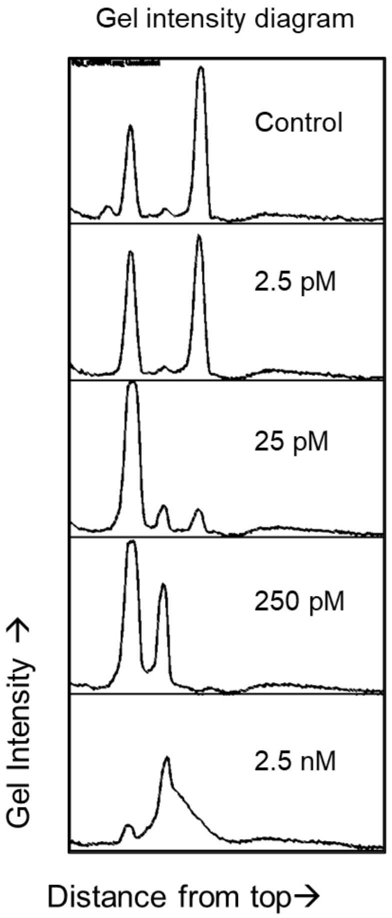


**Fig. S1:** Intensity plot of rCRISPR assay (gel image shown in Fig. 2d). Abbreviations: pM, picomolar; nM, nanomolar.

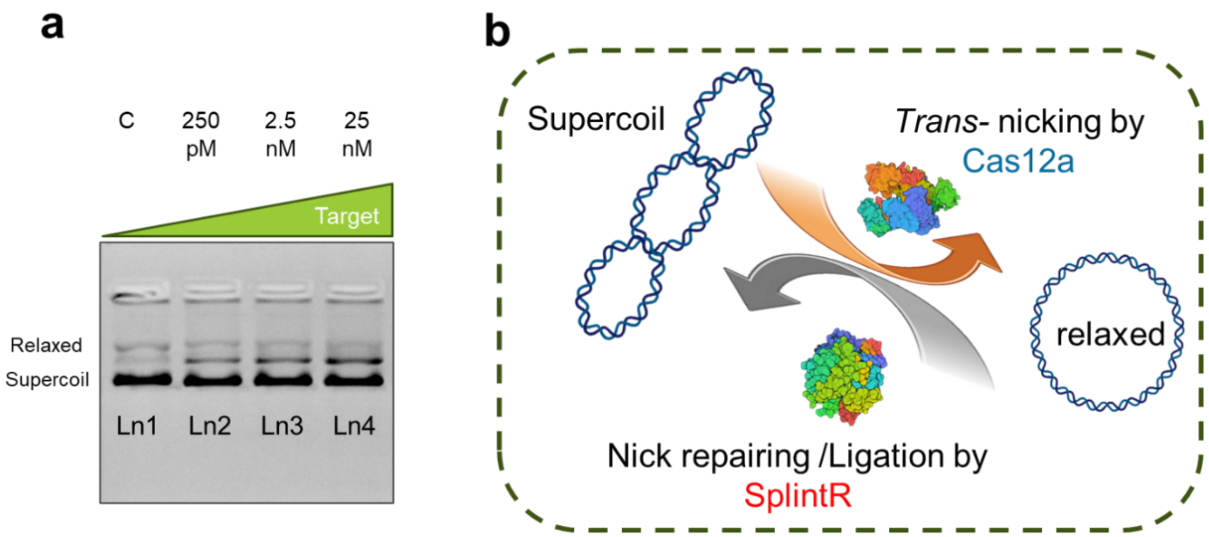


**Fig. S2:** RCA-rCRISPR assay for detecting RNA target. (a) Gel electrophoresis (1% agarose gel and 1×TBE buffer) results demonstrating 2-hr RCA initiated CRISPR-Cas12a induced trans-nicking of phix174 for various target concentrations. (b) Schematic showing how SplintR could revert DNA supercoil relaxation. Abbreviations: c, negative control; pM, picomolar; nM, nanomolar; Ln, lane; TBE, Tris-borate EDTA.

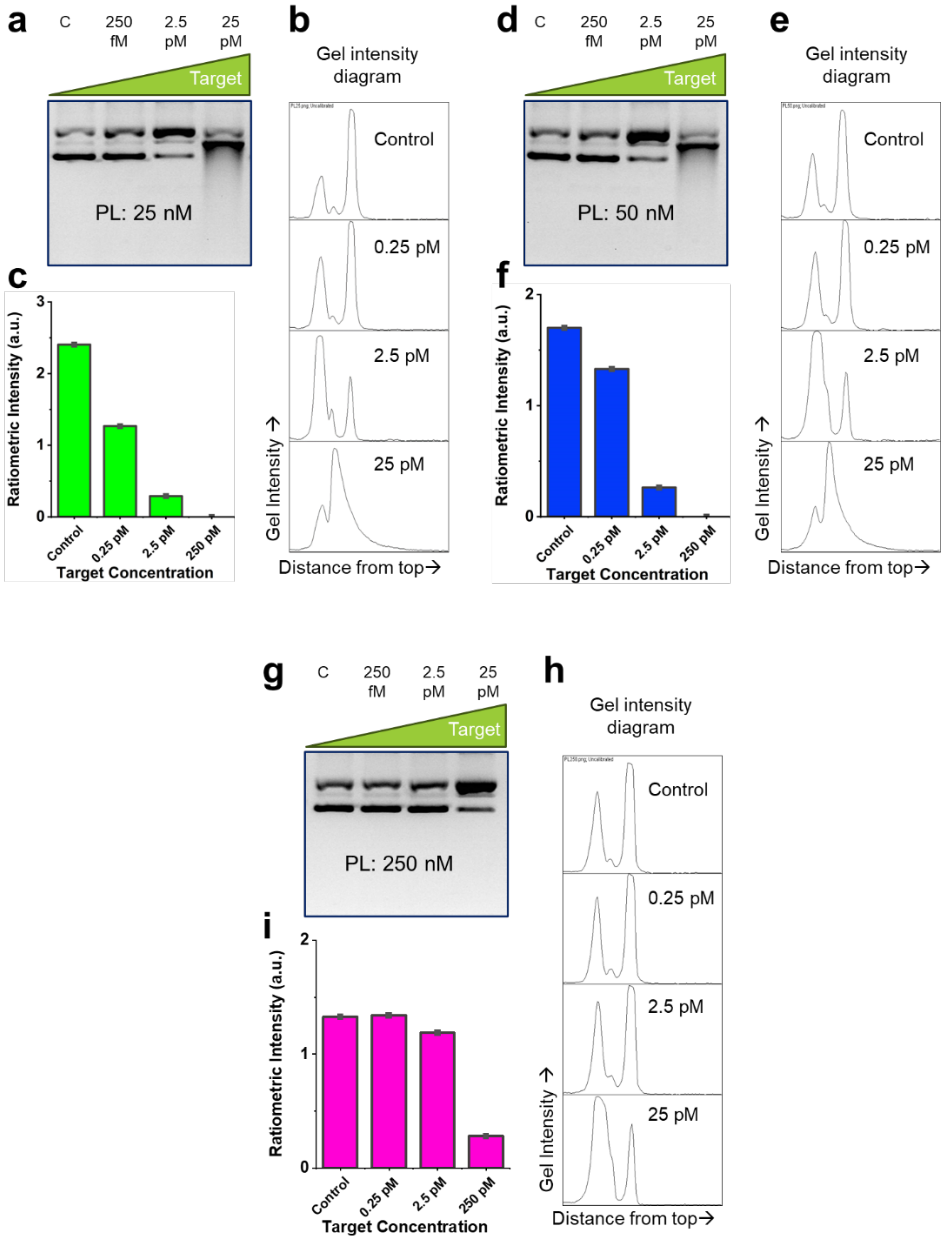


**Fig. S3:** Effect of Padlock concentration. Gel electrophoresis (1% agarose gel and 1×TBE buffer) results demonstrating RNA detection using rCRISPR assay with (a) 25 nM (d) 50 nM, and (g) 250 nM padlock. (b, e, h) Intensity plot diagrams for each lane of the gel in a, d, and g. Bar chart showing assay results with (c) 25 nM (f) 50 nM, and (i) 250 nM padlock. Abbreviations: PL, padlock; c, negative control; pM, picomolar; fM, femtomolar; Ln, lane; TBE, Tris-borate EDTA.

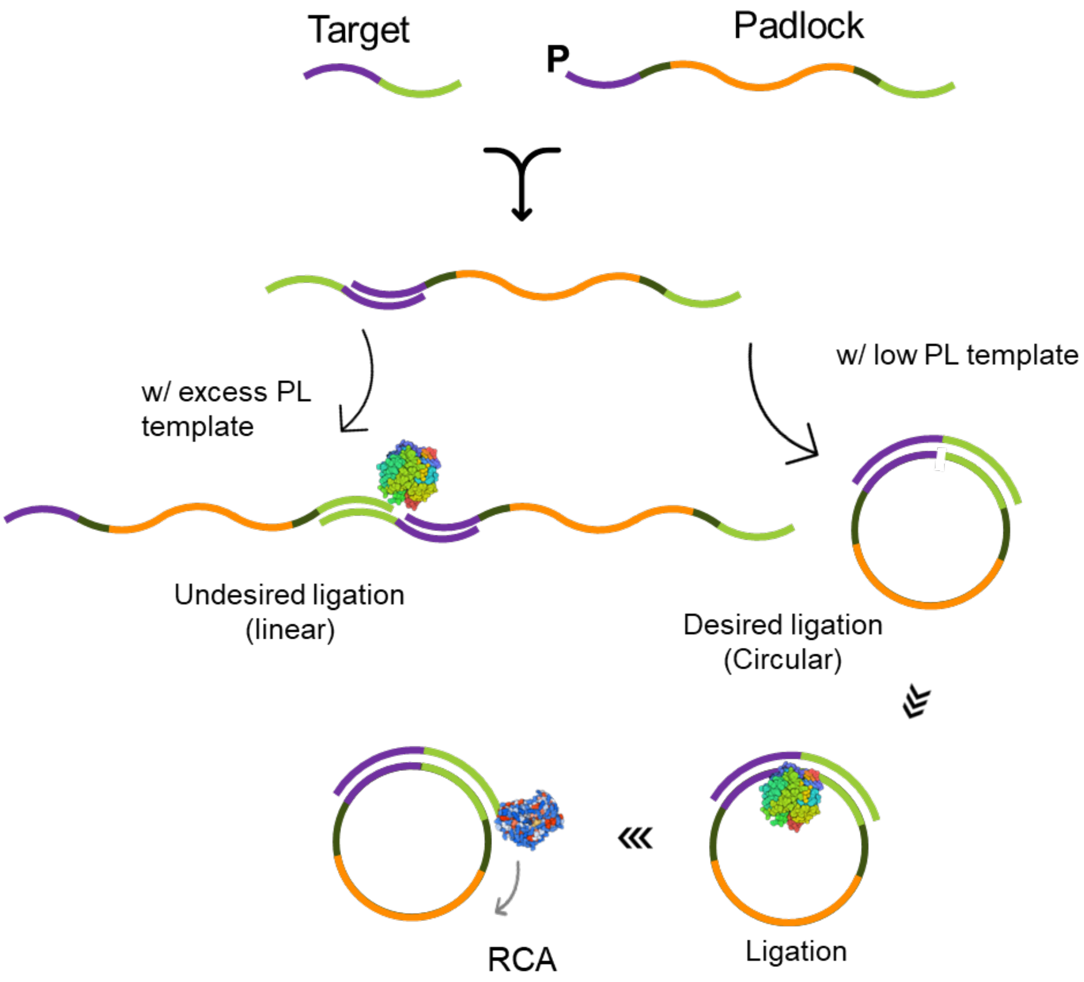


**Fig. S4:** Possible mechanism of how higher padlock concentration inhibit RCA reaction. Abbreviations: PL, padlock; RCA, rolling circle amplification.

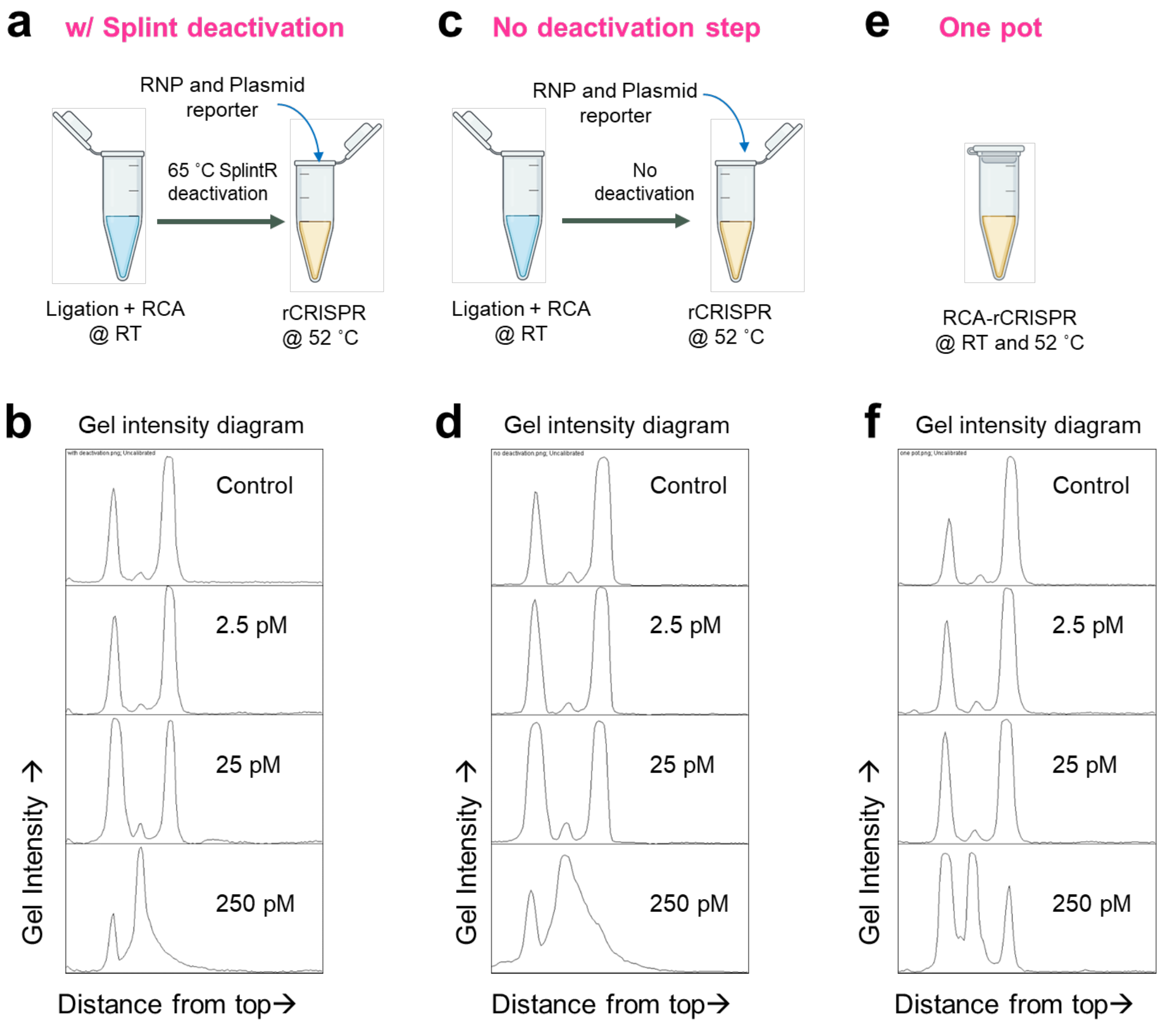


**Fig. S5:** Assay integration. Schematic showing experimental procedure (a) with ligase deactivation (c) without ligase deactivation, and (e) of one pot assay. (b, d, f) Intensity plot diagrams for each lane of the gel in Fig. 4a, 4d, and 4g. Abbreviations: pM, picomolar; RCA, rolling circle amplification.

### HIV detection using the rCRISPR technique

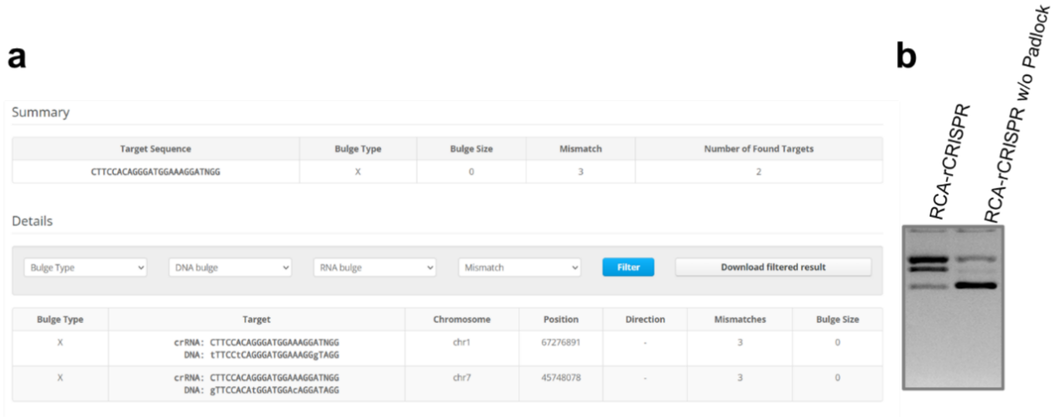


**Fig. S6:** Evaluation of false positives. (a) Off-target checking tool: Cas-OFFinder. (b) Nicking endonuclease and DNase contamination evaluation. Abbreviations: RCA, rolling circle amplification.

### Towards POC detection (equipment-free detection)

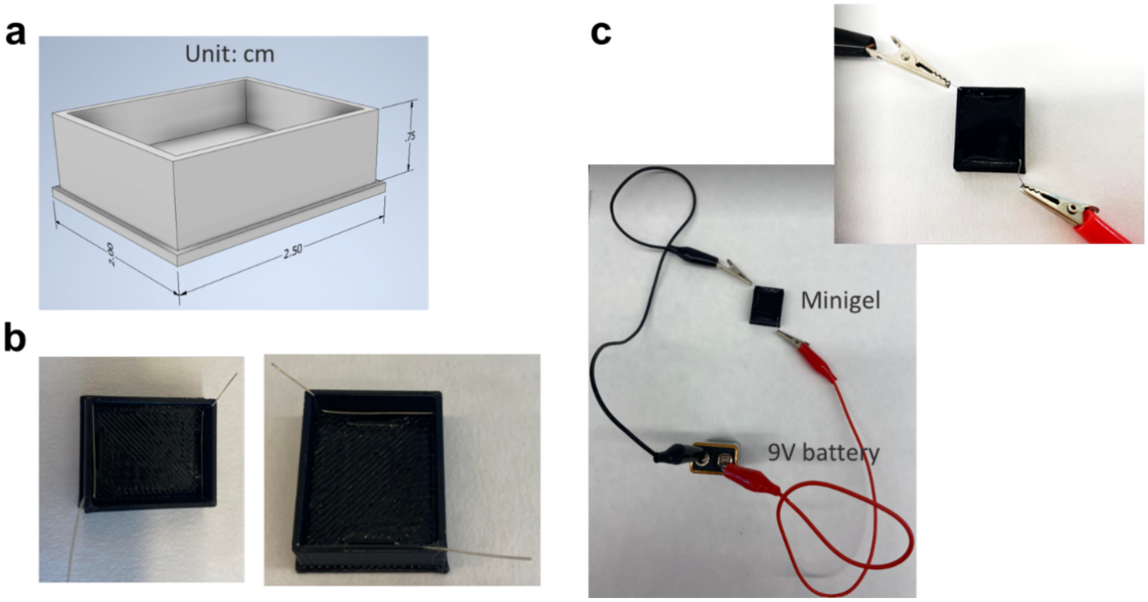


**Fig. S7:** Fabrication of minigel cassette. (a) AutoCAD drawing of minigel cassete. (b) Installation of electrodes using Pt wire. (c) Electrodes connection with a 9v battery.

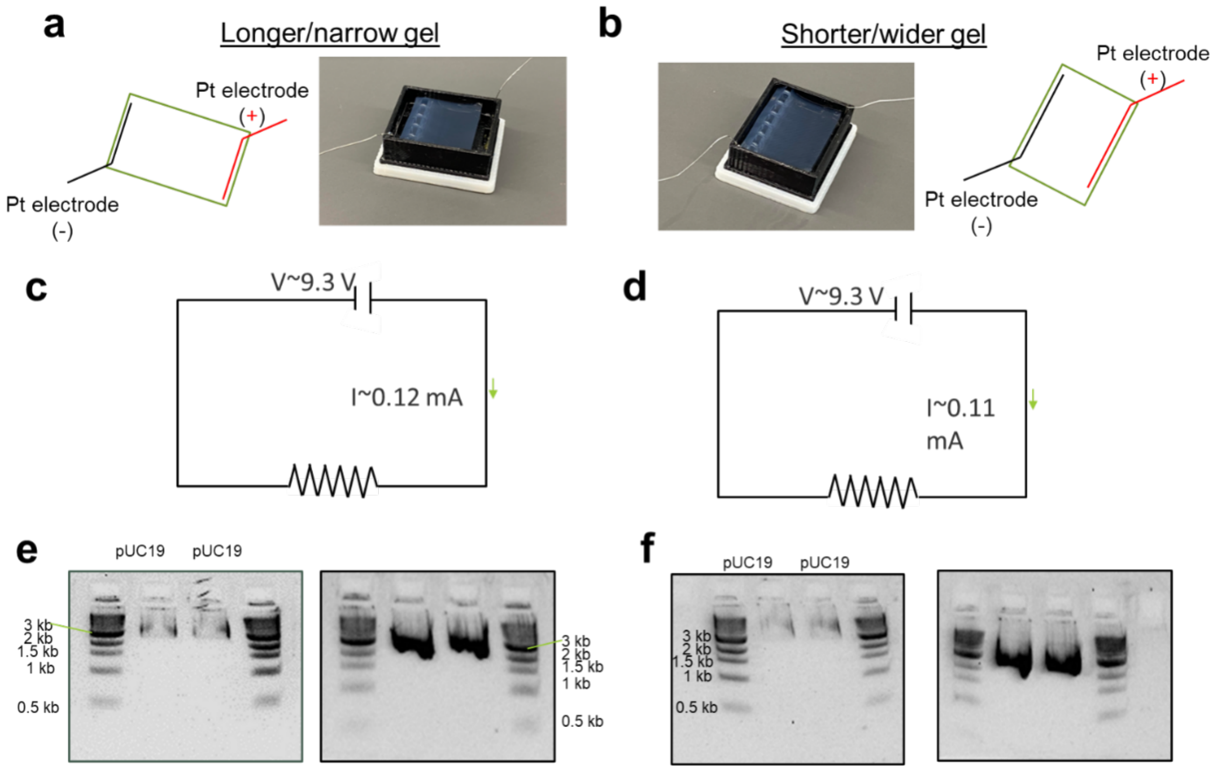


**Fig. S8:** Operational testing of mini-gel using 9v battery. (a,c,e) Longer/narrow gel. (b,d,f) Shorter/wider gel. Abbreviations: Pt, platinum; V, voltage, I, current; kb, kilobase pair.

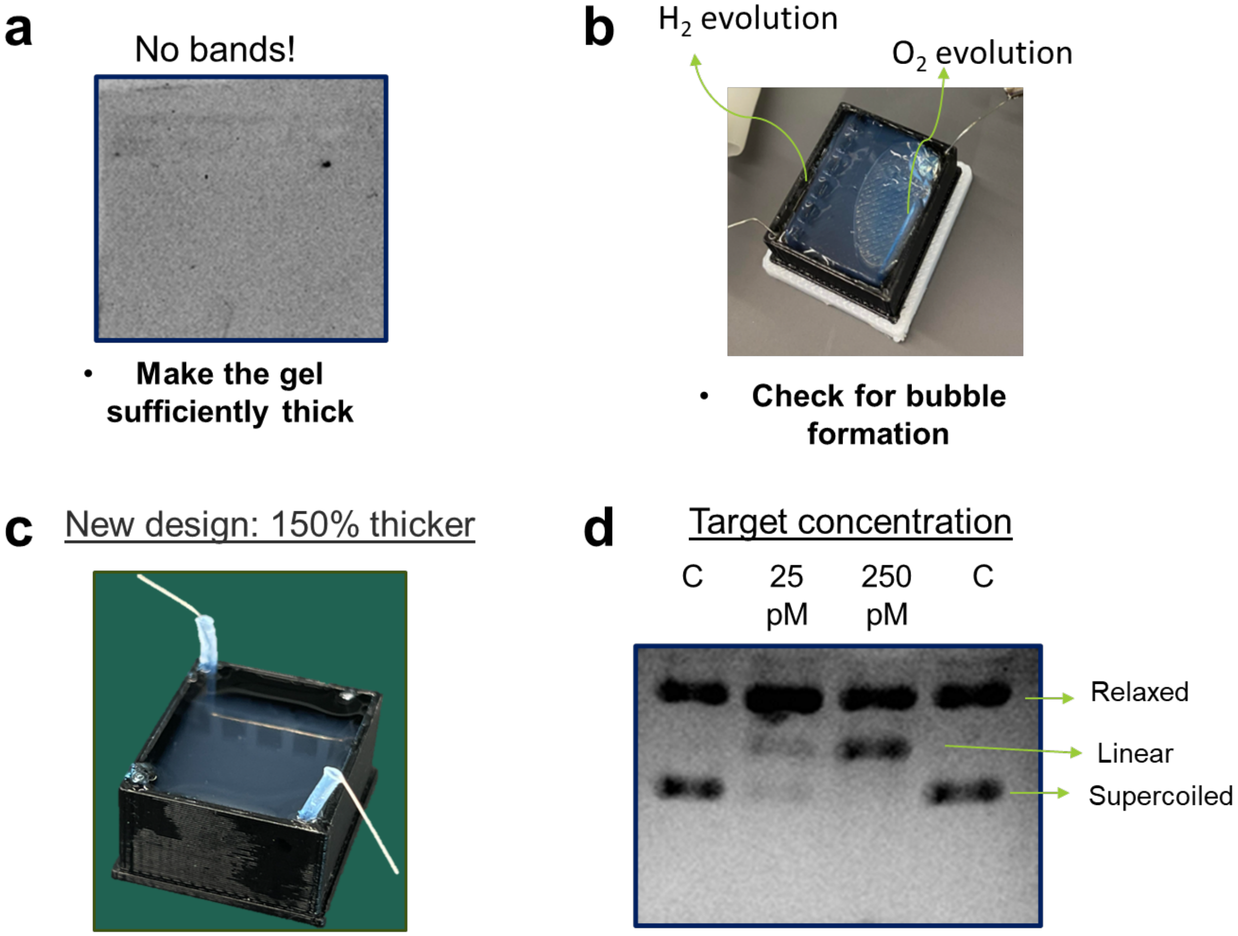


**Fig. S9:** System modifications. (a) Thinner gel resulted in no DNA band. (b) H2/O2 evolution pushed the gel to float. (c) New design that allows thicker gel. (d) phix174 DNA band detection after ratiometric CRISPR assay. Abbreviations: C, control; pM, picomolar.

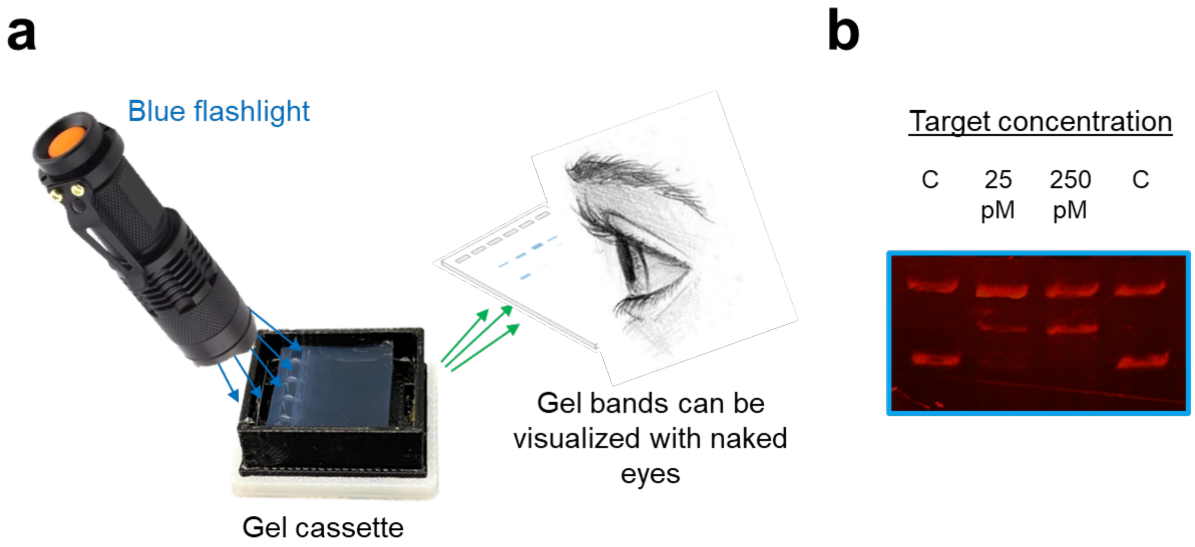


**Fig. S10:** Gel visualization (without transilluminator). (a) Naked eye minigel DNA band detection using a blue light torch. (b) phix174 DNA band detection after ratiometric CRISPR assay. Abbreviations: C, control; pM, picomolar.

### Universal detection (DNA mutation)

**a** Mutated Gene

•

250bp flanking variant base chr16:8296284

•>CATATTTTTAAAAACATATTCATTAACTTTAAGAGGAAATATGAAGTACTATACTTTTTAAAGAAATATCAATTT
TACCAAAAAAAATCATATTATAGAATTACAGAAATTAGATCTCTTACCTAACATTTTTGTAATGCTTGCTTTGCTAG
GAAAATGAGATCTATTGTTTTCCTTTACTTACTACACCTCAGATATTTTTCTT
CATGAAGACCTCACAGTAAAAATAGGTGATTTTGGTCTAGCCACAGAGAAATCTCGATGGAGTGGGTCCCATC
AGTTTGAACAGT
TGTCTGGATCCATTTTGTGGATGGTAAGAATTGAGGCCATTTCTCCATTAATTAAATTTTTGGACCCTGAGGTG
CTACTGAGTGACTAGAAAATCTTTGAAGGTTTCGACTAGTATTTTCATAATTCCCAGTATTCACAAAAATCAATG
TTGATCTTATTTTTTATGTAAATAAAATTTAACTTTTTTCTTTATCCTTAAAAAGAGTATTAT

**b** Wt gene

•

250bp flanking variant base chr16:8296284

•>CATATTTTTAAAAACATATTCATTAACTTTAAGAGGAAATATGAAGTACTATACTTTTTAAAGAAATATCAATT
TTACCAAAAAAAATCATATTATAGAATTACAGAAATTAGATCTCTTACCTAACATTTTTGTAATGCTTGCTTTGCT
AGGAAAATGAGATCTATTGTTTTCCTTTACTTACTACACCTCAGATATTTTTCTT
CATGAAGACCTCACAGTAAAAATAGGTGATTTTGGTCTAGCCACAG**T**GAAATCTCGATGGAGTGGGTCCCAT
CAGTTTGAACAGT
TGTCTGGATCCATTTTGTGGATGGTAAGAATTGAGGCCATTTCTCCATTAATTAAATTTTTGGACCCTGAGGT
GCTACTGAGTGACTAGAAAATCTTTGAAGGTTTCGACTAGTATTTTCATAATTCCCAGTATTCACAAAAATCAA
TGTTGATCTTATTTTTTATGTAAATAAAATTTAACTTTTTTCTTTATCCTTAAAAAGAGTATTAT

**Fig. S11:** SNP location of the canine *BRAF* mutation. (a) Mutated gene. (b) Wild type gene.

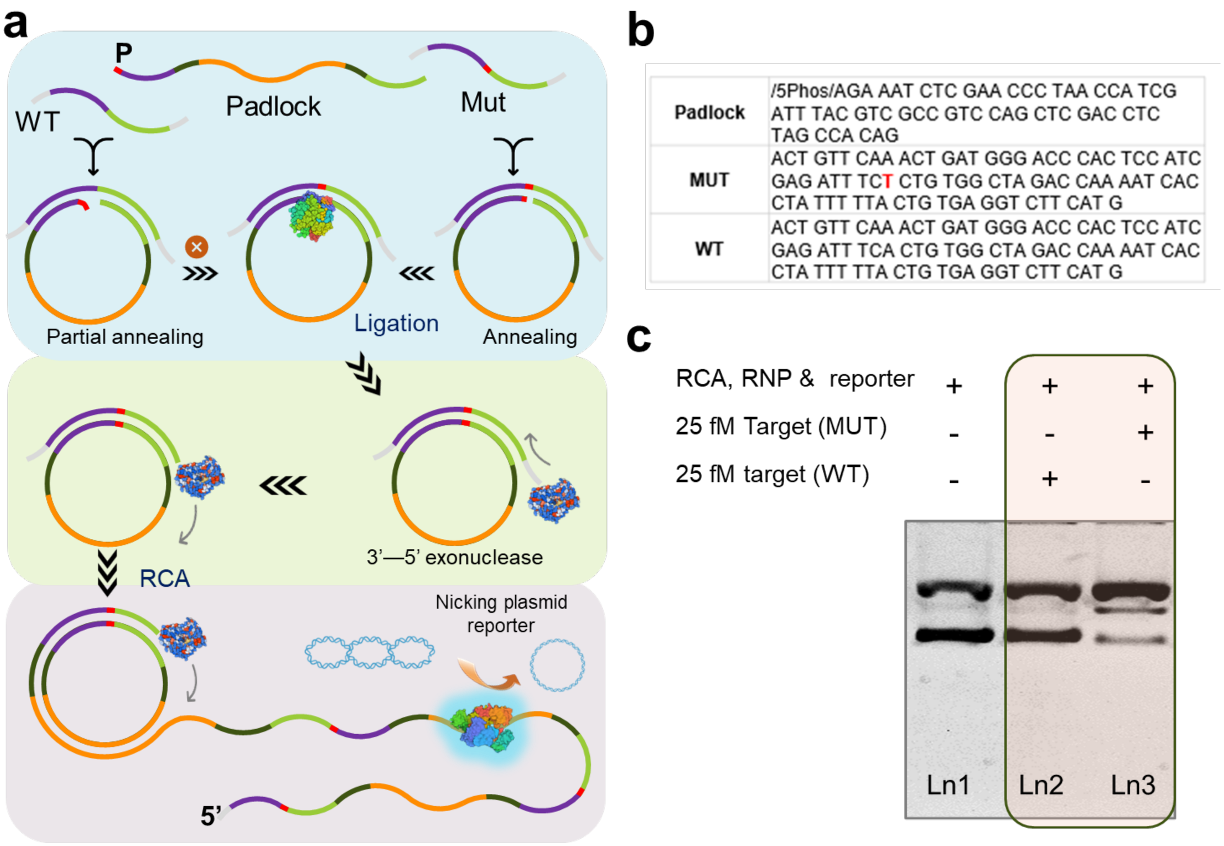


| | |
|---|---|
| Padlock | /5Phos/AGA AAT CTC GAA CCC TAA CCA TCG ATT TAC GTC GCC GTC CAG CTC GAC CTC TAG CCA CAG |
| MUT | ACT GTT CAA ACT GAT GGG ACC CAC TCC ATC GAG ATT TCT CTG TGG CTA GAC CAA AAT CAC CTA TTT TTA CTG TGA GGT CTT CAT G |
| WT | ACT GTT CAA ACT GAT GGG ACC CAC TCC ATC GAG ATT TCA CTG TGG CTA GAC CAA AAT CAC CTA TTT TTA CTG TGA GGT CTT CAT G |

| | Ln1 | Ln2 | Ln3 |
|---|---|---|---|
| RCA, RNP & reporter | + | + | + |
| 25 fM Target (MUT) | - | - | + |
| 25 fM target (WT) | - | + | - |

**Fig. S12**: SNP detection using RCA-rCRISPR strategy. (a) Assay mechanism showing how SNP can be detected with the RCA-initiated rCRISPR technique. (b) DNA sequences (red colored 'T' is the point mutation) used for detection. (c) Gel electrophoresis (1% agarose gel and 1×TBE buffer) results demonstrating the rCRISPR assay result for SNP detection. Abbreviations: RCA, rolling circle amplification; RNP, ribonucleoprotein; WT, wild type; MUT, mutation; Ln, lane; fM, femtomolar; TBE, Tris-borate EDTA.

**Table S1. Oligos used for the study.**

| Item | Sequence (5'-3') |
|---|---|
| gRNA | UAAUUUCUACUAAGUGUAGAUCGUCGCCGUCCAGCUCGACC |
| Amplicon (ssDNA) | GGT CGA GCT GGA CGG CGA CG |
| Padlock_RNA | /5Phos/CTG ATA AGC TAA GAT ACC CTA ACC ATC GAT TTA CGT CGC CGT CCA GCT CGA CCT CAA CAT CAG T |
| Padlock_HIV | /5Phos/CCC TGT GGA AGA CCC TAA CCA TCG ATT TAC GTC GCC GTC CAG CTC GAC CAT CCT TTC CAT |
| Padlock_dogMut | /5Phos/AGA AAT CTC GAA CCC TAA CCA TCG ATT TAC GTC GCC GTC CAG CTC GAC CTC TAG CCA CAG |
| RNA | UAGCUUAUCAGACUGAUGUUGA |
| HIV synthetic target | GUGCUUCCACAGGGAUGGAAAGGAUCAC |
| DogMut | ACT GTT CAA ACT GAT GGG ACC CAC TCC ATC GAG ATT TCT CTG TGG CTA GAC CAA AAT CAC CTA TTT TTA CTG TGA GGT CTT CAT G |
| DogWT | ACT GTT CAA ACT GAT GGG ACC CAC TCC ATC GAG ATT TCA CTG TGG CTA GAC CAA AAT CAC CTA TTT TTA CTG TGA GGT CTT CAT G |
